# Evolution of functional diversity in the holozoan tyrosine kinome

**DOI:** 10.1101/2021.08.03.454916

**Authors:** Wayland Yeung, Annie Kwon, Rahil Taujale, Claire Bunn, Aarya Venkat, Natarajan Kannan

## Abstract

The emergence of multicellularity is strongly correlated with the expansion of tyrosine kinases, a conserved family of signaling enzymes that regulates pathways essential for cell-to-cell communication. Although tyrosine kinases have been classified from several model organisms, a molecular-level understanding of tyrosine kinase evolution across all holozoans is currently lacking. Using a hierarchical sequence constraint-based classification of diverse holozoan tyrosine kinases, we construct a new phylogenetic tree that identifies two ancient clades of cytoplasmic and receptor tyrosine kinases separated by the presence of an extended insert segment in the kinase domain connecting the D and E helices. Present in nearly all receptor tyrosine kinases, this fast-evolving insertion imparts diverse functionalities such as post-translational modification sites and regulatory interactions. Eph and EGFR receptor tyrosine kinases are two exceptions which lack this insert, each forming an independent lineage characterized by unique functional features. We also identify common constraints shared across multiple tyrosine kinase families which warrant the designation of three new subgroups: Src Module (SrcM), Insulin Receptor Kinase-Like (IRKL), and Fibroblast, Platelet-derived, Vascular, and growth factor Receptors (FPVR). Subgroup-specific constraints reflect shared autoinhibitory interactions involved in kinase conformational regulation. Conservation analyses describe how diverse tyrosine kinase signaling functions arose through the addition of family-specific motifs upon subgroup-specific features and co-evolving protein domains. We propose the oldest tyrosine kinases, IRKL, SrcM, and Csk, originated from unicellular pre-metazoans and were co-opted for complex multicellular functions. The increased frequency of oncogenic variants in more recent tyrosine kinases suggests that lineage-specific functionalities are selectively altered in human cancers.

## Introduction

Tyrosine kinases propagate cellular signals through the phosphorylation of tyrosine residues on protein substrates. Forming a monophyletic group within the larger protein kinase superfamily, tyrosine kinases diverged from serine-threonine kinases prior to the emergence of opisthokonts (animals and fungi) (Suga et al. 2012; Hunter 2014), which are estimated to be over a billion years old (Parfrey et al. 2011). While their detection in unicellular pre-metazoans such as choanoflagellates and filastereans has indicated the fundamental roles of tyrosine kinases in the evolution of multicellularity (King et al. 2008; Miller 2012; Tong et al. 2017), their subsequent expansion throughout metazoan evolution is associated with the evolution of diverse metazoan body plans and complex biological systems such as the nervous, vascular, and immune systems (Liu et al. 2011; Miller 2012). Given the vast diversity of tyrosine kinases, the diversification events that gave rise to the functional repertoire of tyrosine kinases and the evolutionary timeline of such events has not been fully explored.

A classification of the protein kinome into evolutionarily and functionally related families (here on referred to as the KinBase classification) was achieved two decades ago following the sequencing and comparative genomic analyses of model organism genomes including human (G. Manning et al. 2002), mouse (Caenepeel et al. 2004), sea urchin (Bradham et al. 2006), fly (Gerard Manning et al. 2002), nematode (Plowman et al. 1999), sponge (Srivastava et al. 2010), choanoflagellate (King et al. 2008), and yeast (Gerard Manning et al. 2002). In addition, tyrosine kinases can be broadly classified as cytoplasmic or receptor tyrosine kinases based on the presence of transmembrane and extracellular ligand binding domains; however, unlike kinase groups and families defined in the KinBase classification, cytoplasmic and receptor tyrosine kinases do not form monophyletic clades in the kinome tree since receptor tyrosine kinases are believed to have independently emerged multiple times throughout tyrosine kinase evolution (Robinson et al. 2000; Suga et al. 2012). The KinBase classification has subsequently become a foundation for comparative studies to study the conservation and divergence of kinase sequence, structure, and function. For example, previous studies of the patterns of sequence conservation and variation across kinase families and groups have provided important insights into the unique regulatory spine of tyrosine kinases relative to serine/threonine kinases (Oruganty et al. 2013; Mohanty et al. 2016), as well as into regulatory mechanisms that evolved uniquely in the EGFR (Mirza et al. 2010), Eph (Kwon et al. 2018), and Tec (Amatya et al. 2019) families of tyrosine kinases.

In addition to the uniquely evolved features across different tyrosine kinase families, similarities across some tyrosine kinase families have also been noted. For example, the recently termed “Src module”, which consists of a tyrosine kinase domain and N-terminal SH3 and SH2 domains, is found across the Src, Abl, Tec, and Csk families, and structural and solution studies have determined that a similar autoinhibitory configuration of the Src module is shared across members of the Src, Abl, and Tec families (Shah et al. 2018). Because previous classifications of protein kinases were determined by analyzing branching points in phylogenetic trees of diverse kinase domain sequences, along with analysis of common domain structures and known biological functions, the existence of evolutionarily related and functionally relevant higher order groupings of families within the kinase classification has not yet been systematically explored.

Here, we determine a novel hierarchical, constraint-based classification of the tyrosine kinome that newly identifies three evolutionary subgroupings of tyrosine kinase families based on the selective conservation of sequence motifs in the kinase domain, which encode common autoinhibitory conformations. In addition, we illustrate an evolutionary timeline of how unique kinase functions have expanded on shared subgroup-specific features through duplication events, evolutionary selection of family-specific motifs, and domain shuffling to give rise to the vast repertoire of tyrosine kinase signaling observed throughout metazoans. A closer examination of tyrosine kinase phylogeny in light of constraint-based tyrosine kinase subgroups reveals new insights into the evolutionary conservation or divergence of subgroups, as well as the unique signaling features that may have emerged from three separate monophyletic clades of receptor tyrosine kinases. In particular, we note the early emergence of two major clades of holozoan tyrosine kinases distinguished by the presence (or absence) of an insert between the αD and αE helices of the kinase domain, where tyrosine kinases containing the insert comprise the majority of metazoan receptor tyrosine kinases. Our classification of the tyrosine kinome and the approach used in this study set a new precedent for the classification and evolutionary study of protein kinases and other large protein families.

## Results

### A hierarchical, constraint-based classification of the holozoan tyrosine kinome reveals new tyrosine kinase subgroups

To generate a comprehensive classification of the holozoan tyrosine kinome, we generated a multiple sequence alignment of 44,639 tyrosine kinase sequences spanning 586 species (see methods for details). We then used a Bayesian Partitioning with Pattern Selection (BPPS) algorithm to classify aligned sequences into hierarchical clusters based on the patterns of conservation and variation in aligned tyrosine kinase domain sequences **(Figure 1A)** (Neuwald 2011; Neuwald 2014). Each cluster is distinguished by co-occurring sequence motifs which are highly conserved within the cluster, but strikingly different outside of the cluster. By sampling different clustering hierarchies and highly distinguishing sequence motifs, we defined an optimal hierarchy for holozoan tyrosine kinases based on the log-probability ratio (LPR) scores, which quantifies the contribution of conserved sequence patterns to the classification/clustering measured in natural units of information (nats) (Neuwald 2011). The total LPR score for the optimized holozoan tyrosine kinome classification was 459825.35 nats, which is higher than the LPR score for KinBase classification of tyrosine kinases (443759.95 nats) **(Supplementary Figure S1)**.

**Figure 1:**
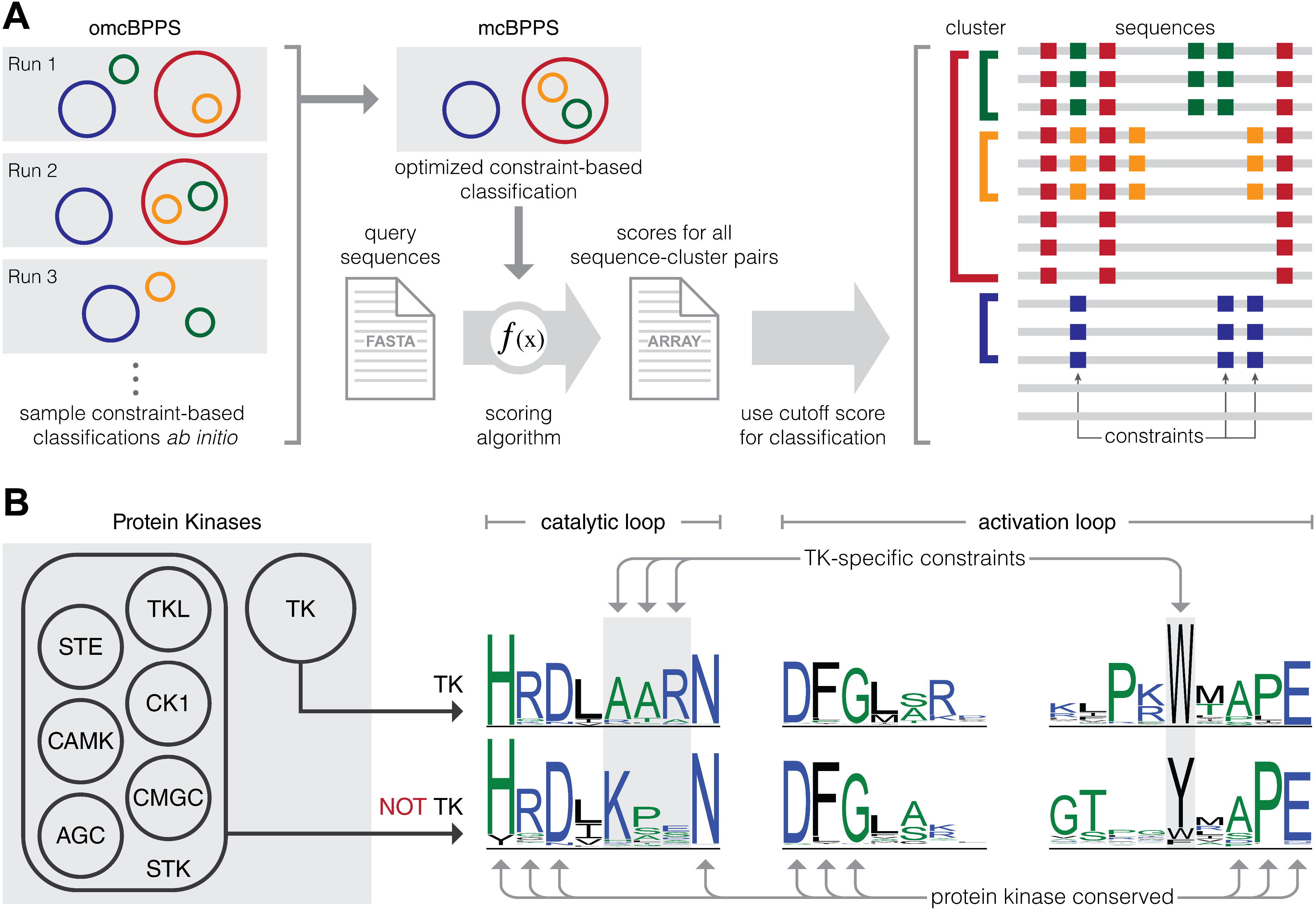
Workflow for constraint-based hierarchical clustering of tyrosine kinase sequences. (A) Multiple omcBPPS runs were used to sample various constraint-based hierarchical classifications for tyrosine kinases. An optimal constraint-based classification was determined based on LPR scores calculated using mcBPPS. Next, a scoring algorithm was used to score sequence-cluster pairs and to include or exclude tyrosine kinase sequences from each cluster defined in the optimal classification. An example constraint-based hierarchical classification is shown on the right. Clusters are represented as colored brackets, sequences are represented as grey lines, and constraints specific to each cluster are represented by squares colored according to the cluster to which they belong. For example, sequences in the green cluster share sequence constraints denoted by green squares, which are not found in sequences outside of the green cluster, as well as sequence constraints denoted by red squares, which are not found in sequences outside of the red cluster. The last two sequences are not included in any cluster as they lack any of the cluster-specific constraints defined in the constraint-based classification. (B) A visual representation of cluster-specific constraints is shown using previously published data on tyrosine kinase-specific constraints (Mohanty et al. 2016). Seven clusters of protein kinases are shown on the left, where tyrosine kinases are clustered separately from other protein kinases. A sequence logo of tyrosine kinase sequences is shown alongside a sequence logo of all other protein kinases. Sequence motifs such as HRD, DFG, and APE are conserved throughout all protein kinases, whereas the catalytic loop AAR motif and the activation loop tryptophan are defined as TK-specific constraints because their conservation is specific to the TK cluster.

Next, we re-classified 34,954 tyrosine kinase sequences from the UniProt reference proteomes database into the optimized hierarchy by quantifying the extent to which individual sequences match cluster-specific motifs (see methods for details). We define this re-classification as a constraint-based classification because this post-processing step eliminates spurious or divergent sequences from clusters that do not score over an optimal cutoff score due to their lack of cluster-specific patterns. The spurious sequences eliminated from each cluster are categorized within the Unclassified family **(Figure 2)**.

**Figure 2:**
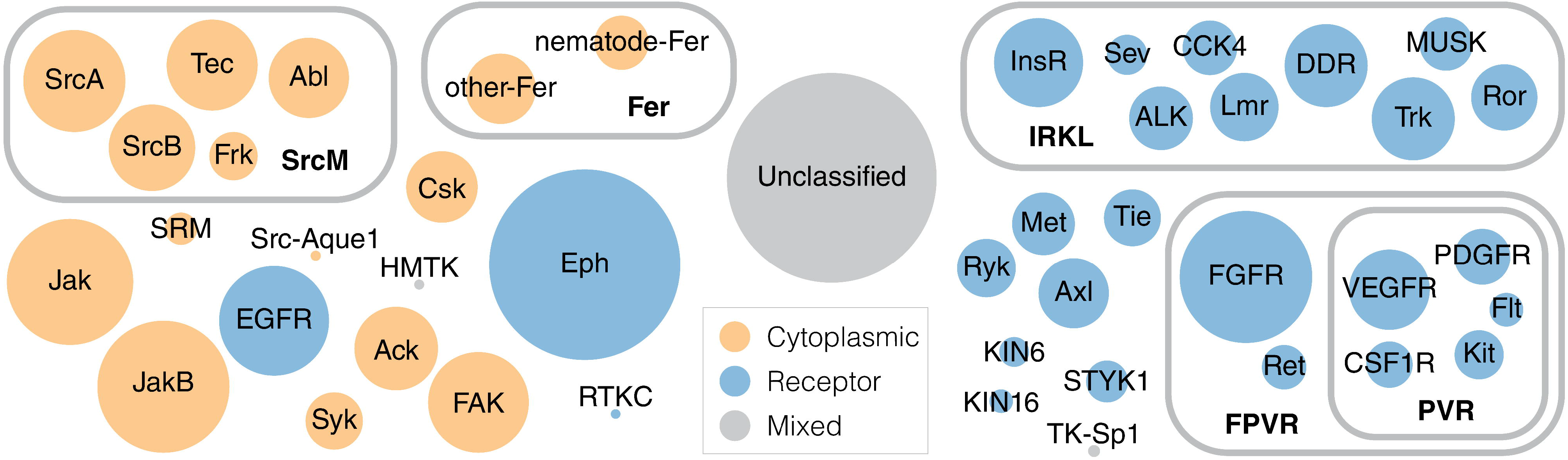
An evolutionary constraint-based hierarchical classification of the tyrosine kinome. The constraint-based hierarchical classification of tyrosine kinases is depicted as a Euler diagram. Each circle represents a distinct cluster of tyrosine kinases and is scaled to the number of sequences in each cluster. Clusters containing cytoplasmic tyrosine kinases are indicated with orange circles, while clusters containing receptor tyrosine kinases are indicated with blue circles. Clusters containing both cytoplasmic and receptor tyrosine kinases are colored grey. For more information, see **Supplemental File S1**.

The new constraint-based hierarchical classification of tyrosine kinases, which is broadly similar to the KinBase classification of the tyrosine kinome, reveals several novel sub-groupings. In particular, the constraint-based classification defines three new subgroupings of tyrosine kinase families which account for nearly half of the tyrosine kinome: the Src Module (SrcM) subgroup, the Insulin Receptor Kinase-Like (IRKL) subgroup, and the Fibroblast, Platelet-derived, and Vascular growth factor Receptors (FPVR) subgroup **(Figure 2)**. The SrcM subgroup differs significantly from the KinBase classification in that it clusters the SrcA, SrcB, and Frk subfamilies of the KinBase-defined Src family within the same subgroup as the Tec and Abl families. Furthermore, SrcM does not include the SRM and sponge-specific Src (Src-Aque1) families. The FPVR subgroup includes seven distinct receptor tyrosine kinase families, where a Platelet-derived, and Vascular growth factor Receptor (PVR) subgroup sub-classifies the VEGFR, PDGFR, Kit, CSF1R, and Flt3 families as a distinct cluster separate from the FGFR and Ret families. Notably, the new classification separates the KinBase PDGFR family into four families (PDGFR, Kit, CSF1R, and Flt3) due to statistically significant sequence constraints that define each of these families, warranting their designation as distinct families. The IRKL subgroup is the largest subgroup, comprising roughly 16% of tyrosine kinase sequences, and encompasses nine receptor tyrosine kinase families, including the insulin receptor kinase family as well as other poorly studied tyrosine kinases such as the CCK4 family of pseudokinases (Jung et al. 2004; Murphy et al. 2013) and the Lmr family which exhibits serine/threonine kinase activity despite its placement into the tyrosine kinase clade (Wang and Brautigan 2002; Ditsiou et al. 2020). We note that some organism-specific tyrosine kinase families defined in the KinBase classification, such as the unique receptor tyrosine kinase families in choanoflagellates (e.g. RTKA, RTKB, etc.) (King et al. 2008; Manning et al. 2008), were not detected due to the limited number of detectable homologs in current sequence databases (see methods).

### Subgroup-specific motifs localize to known autoregulatory sites in the kinase domain

By examining the sequence constraints that define each of the three novel subgroups in light of existing crystal structures, we observe that subgroup-specific motifs are located in known regulatory regions of the kinase domain. For example, the SrcM subgroup conserves a highly distinguishing GxM motif in the β3-αC loop and a GxKF motif in the activation loop that both form important interactions associated with a common Src-like inactive conformation in the activation loop **(Figure 3A)** (Xu et al. 1999; Shah et al. 2018). This Src-like inactive conformation has been observed across diverse SrcM families such as SrcA (Xu et al. 1999), SrcB (Schindler et al. 1999), Abl (Levinson et al. 2006), and Tec (Wang et al. 2015), and similarities between their inactive structures have been previously noted (Shah et al. 2018). That SrcM-specific sequence motifs are located in key regions in the Src-like inactive conformation, suggesting that these residues play key roles the conformational control of SrcM kinase activity. Likewise, the strongest sequence constraints on IRKL tyrosine kinases are associated with a common autoinhibitory conformation of the activation loop **(Figure 3B)**, which has been observed across crystal structures of diverse IRKL members (Hubbard et al. 1994; Artim et al. 2012; Canning et al. 2014; Ditsiou et al. 2020), and is distinct from the autoinhibitory activation loop conformation of SrcM tyrosine kinases. Lastly, the FPVR subgroup of tyrosine kinases is defined by a highly conserved asparagine in the hinge region of the kinase domain **(Figure 3C)**, which engages an autoinhibitory ‘molecular brake’ (Chen et al. 2007) shared across these kinases. Other FPVR-specific sequence motifs are structurally located near the juxtamembrane, which is an important regulatory segment for many receptor tyrosine kinases (Griffith et al. 2004), and may play common structural and functional roles in juxtamembrane mediated regulation across FPVR tyrosine kinases.

**Figure 3:**
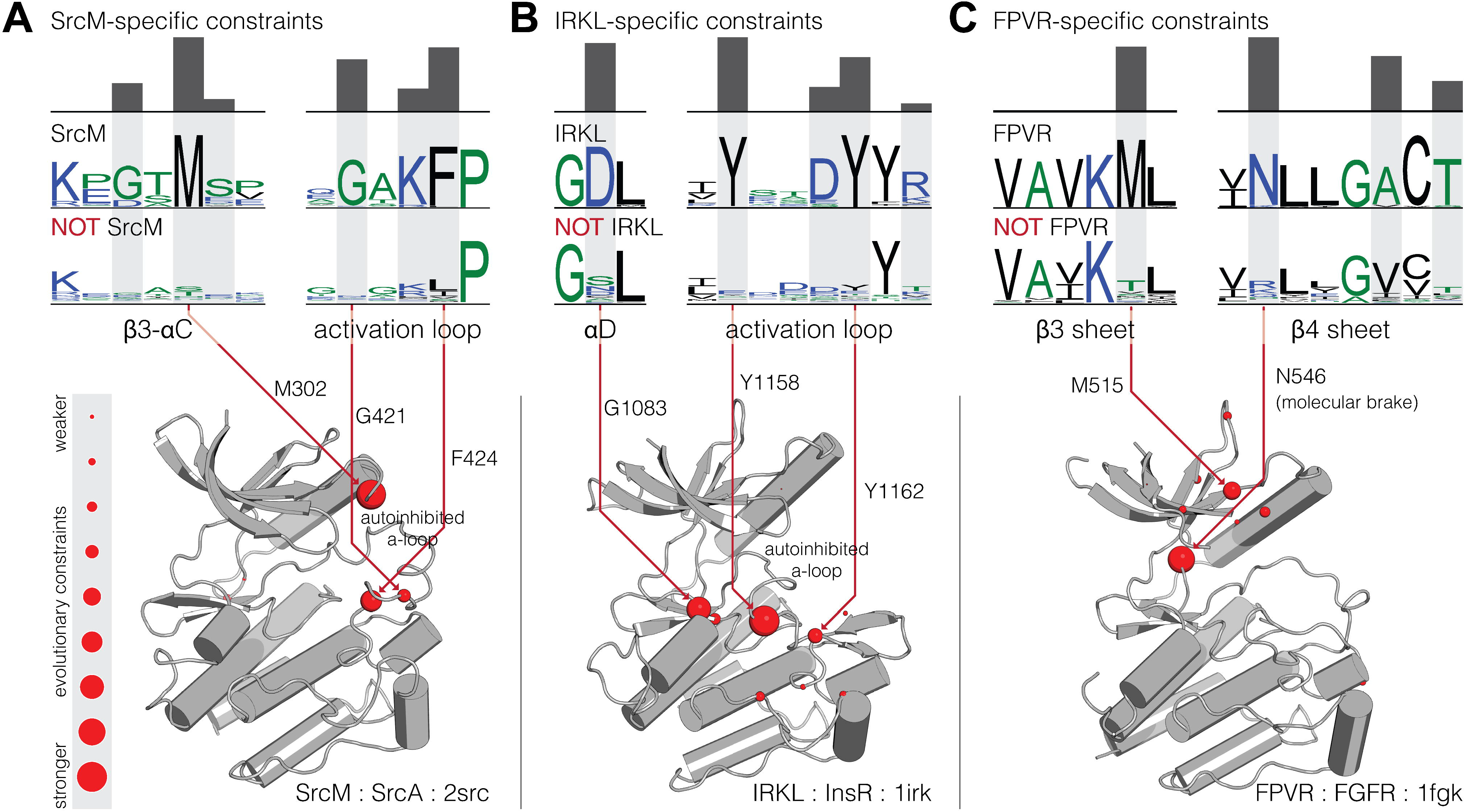
Structural locations of sequence motifs defining the SrcM, IRKL, and FPVR subgroups. Comparative sequence logos and structural mappings of subgroup-specific motifs are shown for the (A) SrcM, (B) IRKL, and (C) FPVR subgroups of tyrosine kinases. Sequence logos for the strongest evolutionary constraints corresponding to each subgroup are shown on top, with comparative sequence logos for sequences outside each subgroup provided below. Evolutionary constraints are highlighted in gray, with the height of the histogram reflecting the degree of divergence at that position between the subgroup and sequences outside the subgroup. Evolutionary constraints are shown as red balls on representative inactive structures of SrcM (Src) (Xu et al. 1999), IRKL (IRK) (Hubbard et al. 1994), and FPVR (FGFR1) (Mohammadi et al. 1996). The size of the red balls represents the strength of the evolutionary constraint. The strongest constraints are labeled with residue numbers corresponding to the position in the representative structures.

### Tyrosine kinase subgroups are anciently conserved across diverse holozoan taxa

In order to infer when each of these tyrosine kinase subgroups and families emerged in evolution, we organized tyrosine kinase subgroups and families based on taxonomic conservation **(Figure 4)**. Tyrosine kinases from the SrcM subgroup, IRKL subgroup, and Csk family are detected across the most diverse holozoan taxa, including in unicellular relatives of metazoans (pre-metazoans) such as filastereans and choanoflagellates. The conservation of these early-emerging tyrosine kinases suggests that they likely played important roles in the evolution of metazoan multicellularity. The FPVR subgroup appears to have emerged later in eumetazoans, following the divergence from early metazoans such as poriferans, which lack the organized tissues observed across eumetazoans. Interestingly, the PVR subgroup within the FPVR subgroup evolved much later in metazoan evolution and can only be detected in deuterostomes. The fact that tyrosine kinases from the SrcM, IRKL, and FPVR subgroups emerged in early stages of metazoan evolution suggests that these tyrosine kinases, their defining sequence motifs, and the regulatory functions associated with the motifs **(Figure 3)** were important in the evolution of metazoan morphologies such as multicellularity and organized tissues. We also note that, as found in previous kinome studies, our constraint-based classification defines several organism-specific tyrosine kinase families, such as the sponge-specific Src-Aque1 family (Srivastava et al. 2010), the nematode specific KIN6 and KIN16 families (Plowman et al. 1999), and the choanoflagellate-specific HMTK and RTKC families (King et al. 2008) **(Figure 4)**.

**Figure 4:**
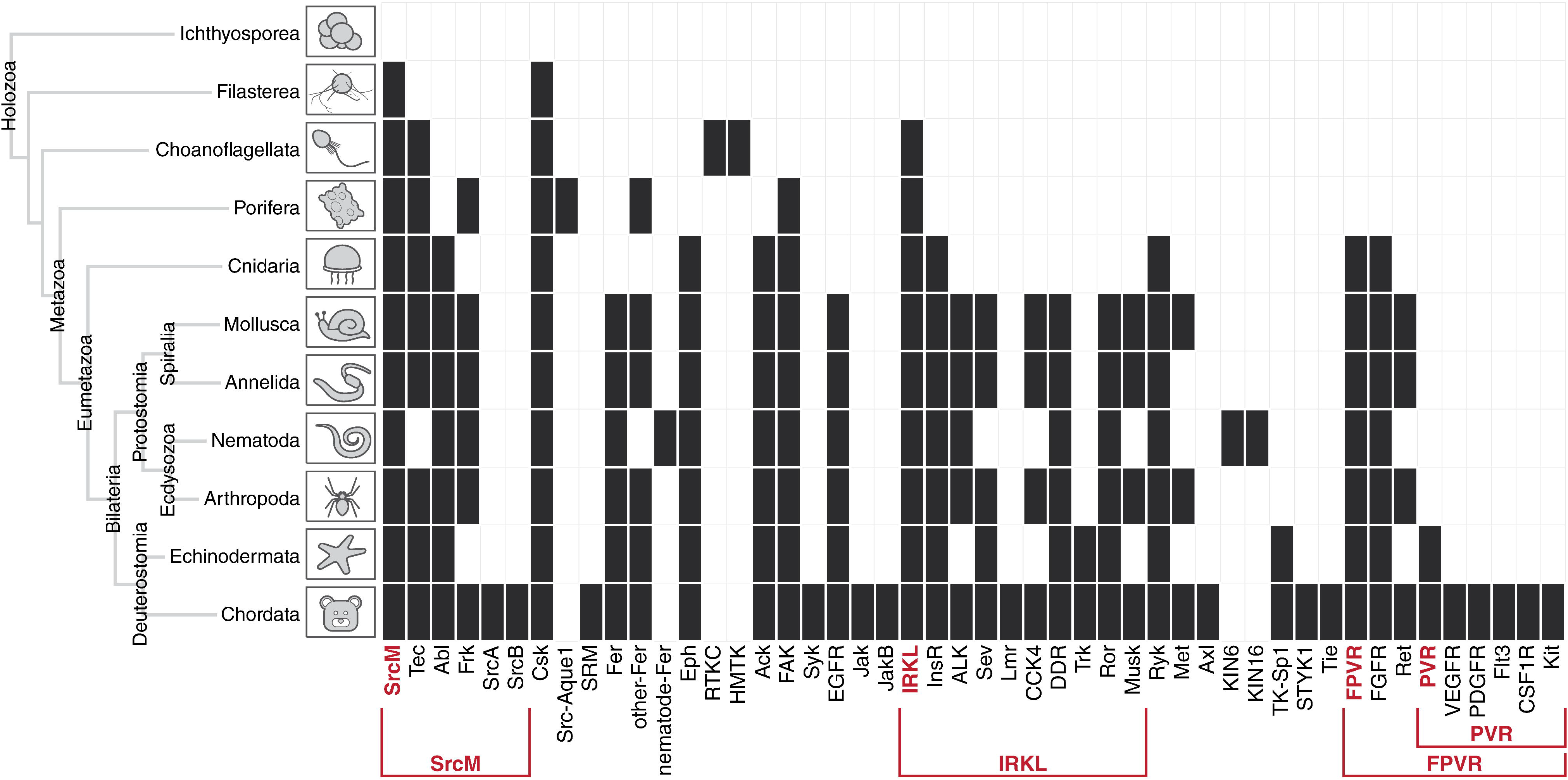
Emergence of tyrosine kinase subgroups and families throughout diverse holozoan taxa. The detection of each tyrosine kinase subgroup and family defined in the constraint-based classification is shown across diverse holozoan taxa, including single-celled relatives of metazoa. Constraint-based tyrosine kinase subgroups and families are shown across the x-axis, and diverse holozoan taxa, and their evolutionary relationships are shown on the y-axis. Cells are marked black if one or more members of a subgroup or family could be detected within a given taxa. For more information, see **Supplemental File S2**.

### Domain shuffling contributed to diverse functions of the SrcM, IRKL, and FPVR kinase domains

In order to further explore the functional diversity of SrcM, IRKL, and FPVR tyrosine kinases, we surveyed the diversity of protein domains present across these subgroups and analyzed their conservation across holozoan taxa **(Figure 5)**. As previously noted, the SrcM tyrosine kinases, as well as the SRM, Csk, and Src-Aque1 families of tyrosine kinases share a core SH3-SH2-kinase domain organization, an anciently conserved (Shah et al. 2018), co-evolving unit (Nars and Vihinen 2001) which can be detected in SrcM orthologs in unicellular pre-metazoans such as choanoflagellates and filastereans **(Figure 5D)**. The Tec family domain architecture, which includes an N-terminal lipid-targeting PH-domain (except in the Tec family member Txk), can be detected in choanoflagellates and filastereans, therefore the SH3-SH2-kinase and PH-SH3-SH2-kinase domain architectures represent the most anciently conserved tyrosine kinase domain structures. Because sequence motifs defining the SrcM subgroup and the Tec family are also anciently conserved **(Figure 4)**, we suggest that these motifs have co-evolved with the SH3-SH2 and PH domains, respectively, and play key functional roles that link the kinase domain with their associated domains. Interestingly, the Abl family domain architecture, which includes a C-terminal F-actin binding domain appended to the SH3-SH2-kinase domain structure, emerged later in metazoan evolution after the divergence of bilaterians from other eumetazoans. However, evolutionary sequence constraints that distinguish the Abl family kinase domain from other SrcM members emerged earlier in metazoan evolution after the emergence of eumetazoans **(Figure 4)**. This suggests that Abl-specific functions of the kinase domain, perhaps substrate-specificity or Abl-specific regulation of catalysis, predated the additional functions imparted by the F-actin binding domain.

**Figure 5:**
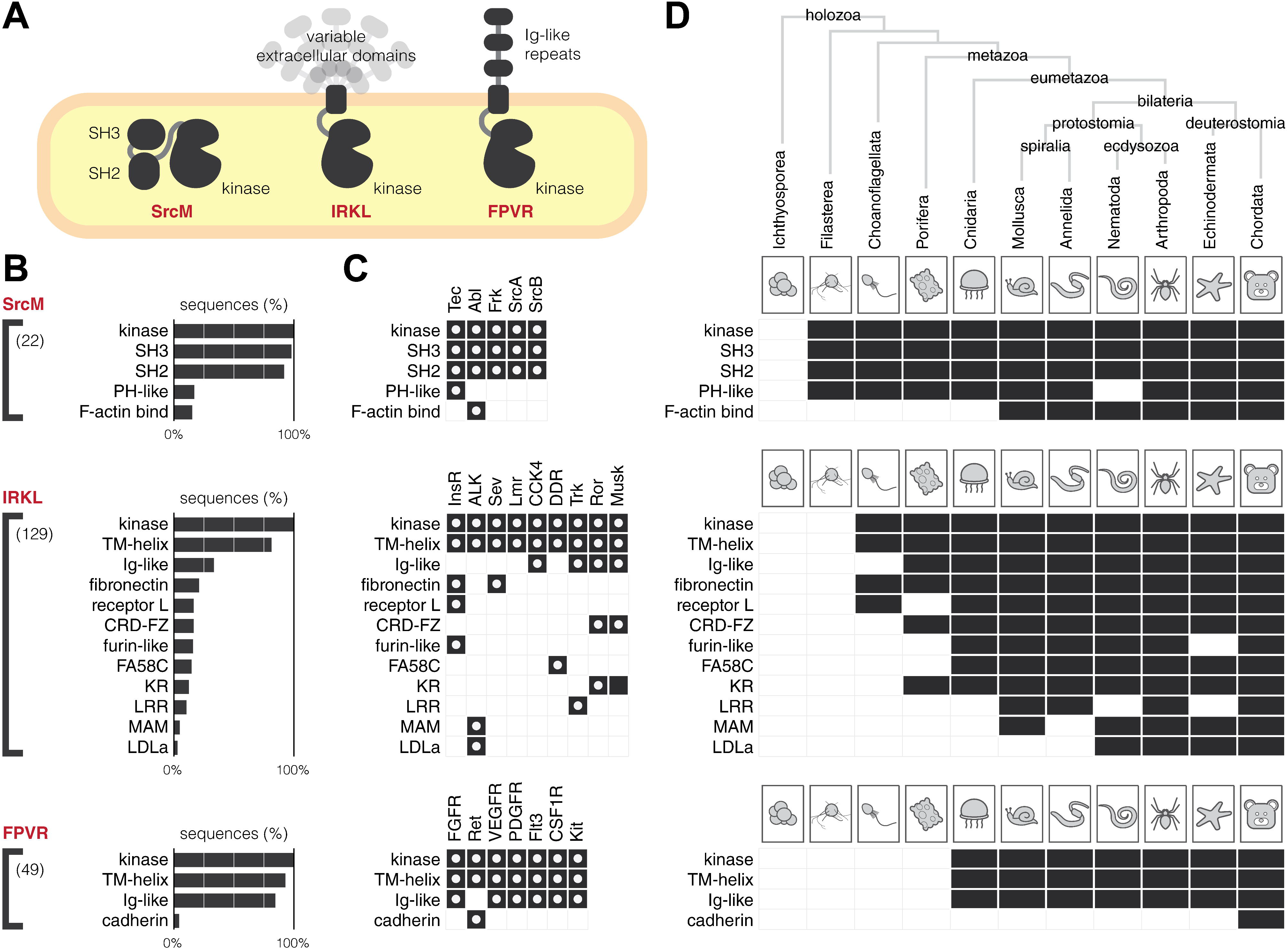
Protein domains associated with SrcM, IRKL, and FPVR tyrosine kinases. (A) A graphical depiction of common protein domain architectures observed across the SrcM, IRKL, and FPVR subgroups. (B) Bar graphs show the frequencies of protein domains detected in SrcM, IRKL, and FPVR sequences. Protein domains that occur in at least 3% of sequences are shown. The value in parentheses denotes the total number of unique protein domains found for sequences of a given subgroup. Consecutive repeat domains have been compressed into a single domain for simplicity. (C) Protein domains found across individual tyrosine kinase families within the SrcM, IRKL, and FPVR subgroups. White dots indicate domains found in human tyrosine kinases in each family. (D) The conservation of protein domains associated with SrcM, IRKL, and FPVR tyrosine kinases across diverse holozoan taxa.

The IRKL and FPVR subgroups encompass the majority of receptor tyrosine kinase families and, despite sharing common kinase domain mechanisms within these subgroups **(Figure 3)**, have diversified functions through the incorporation of various extracellular domains. In fact, the diverse assortment of extracellular domains architectures found throughout the IRKL subgroup may be consistent with extracellular domain shuffling throughout holozoan evolution. Interestingly, the emergence of family-specific extracellular domains often precedes the emergence of family-specific motifs that define receptor tyrosine kinase families. For example, the extracellular fibronectin domain, which is associated with InsR and Sev family receptor tyrosine kinases, can be detected in diverse holozoan taxa, from chordates to choanoflagellates **(Figure 5D)**, however, InsR and Sev-specific sequence motifs in the kinase domain emerged later in metazoan evolution during the emergence of eumetazoans and bilaterians, respectively **(Figure 4A)**. Similarly, Frizzled cysteine-rich (CRD-FZ), Coagulation factor 5/8 C-terminal (FA58C), and leucine-rich repeat (LRR) domains, which are associated with the Ror/Musk, DDR, and Trk families, respectively, emerged before their respective family-specific motifs. The evolutionary emergence of extracellular domains before family-specific motifs in the kinase domain suggests that evolution first diversified extracellular ligand binding before the fine-tuning of family-specific kinase domain functions such as downstream substrate specificity, regulation of catalysis, or intracellular protein-protein interactions. In contrast, in the ALK and Ret families of receptor tyrosine kinases, which uniquely contain LDLa and cadherin extracellular domains, respectively, the emergence of their family-specific sequence motifs predated the addition of their distinctive extracellular domains. These cases suggest that unique family-specific functions in the intracellular portion of receptor tyrosine kinases can also be expanded upon by subsequent shuffling of extracellular domains such that intracellular signaling functions are newly adapted to alternative extracellular ligands. Generally, despite the high degree of conservation of family-specific core domain structures **(Figure 5B, 5C)**, the extreme diversification of SrcM, IRKL, and FPVR kinase domains across holozoans is evident in the huge number of unique protein domains that can be detected across sequences.

### A representative phylogeny of the holozoan tyrosine kinome reveals new insights into tyrosine kinase evolution

To better understand the evolutionary relationships between tyrosine kinase subgroups and families, we constructed a phylogenetic tree using maximum likelihood, which models the natural process of sequence variation and finds a tree that best describes the evolutionary history of diverse protein sequences (see methods for details). By integrating our constraint-based classification of tyrosine kinases with our phylogenetic tree **(Figure 6A)**, we can now infer the evolutionary history of sequence constraints imposed on tyrosine kinase subgroups and families defined in our constraint-based classification. For example, the IRKL, FPVR, and PVR subgroups, which were defined in our constraint-based classification due to their conservation of a set of subgroup-specific motifs, each form monophyletic clades in the phylogenetic tree, demonstrating that all tyrosine kinases within these subgroups have both maintained their respective subgroup-specific motifs and have descended from a common evolutionary ancestor. In contrast, the SrcM subgroup does not form a monophyletic clade in the phylogenetic tree. Instead, families within the SrcM subgroup share a monophyletic clade with the SRM, Src-Aque1, and Csk families, all of which likely descended from a common ancestor which conserved SrcM-specific motifs. However, that the SRM, Src-Aque1, and Csk families were not included in our constraint-based definition of the SrcM subgroup signifies that these families independently diverged from the rest of the SrcM subgroup through variations in the canonical SrcM-specific motifs, as well as accumulating additional variations contributing to functional divergence. Furthermore, examining SRM-specific, Src-Aque1-specific, and Csk-specific motifs unique to each of these families alongside SrcM subgroup-specific motifs will reveal how a common SH3-SH2-kinase domain organization **(Figure 5)** (Shah et al. 2018) has diverged along these various lineages to innovate divergent regulatory functions on a shared domain architecture. Our phylogenetic tree also confirms, along with our constraint-based classification, that the Lmr family belongs within the IRKL subgroup/clade despite detectable serine/threonine activity. Further sequence analysis shows that the Lmr kinases have regained a serine/threonine kinase-specific histidine (LMTK3^His260^) in the αE helix which tyrosine kinases selectively lost upon diverging from the serine/threonine kinases **(Supplemental Figure S5)** (Mohanty et al. 2016). The reemergence of this histidine may explain why Lmr kinases have regained serine/threonine activity. In addition, the evolutionary relationships described by our phylogeny of the holozoan tyrosine kinome are independently support by conserved intron and phase positions in the kinase domain **(Supplemental Figure S6**) (Brunet et al. 2016; Brunet et al. 2017).

**Figure 6:**
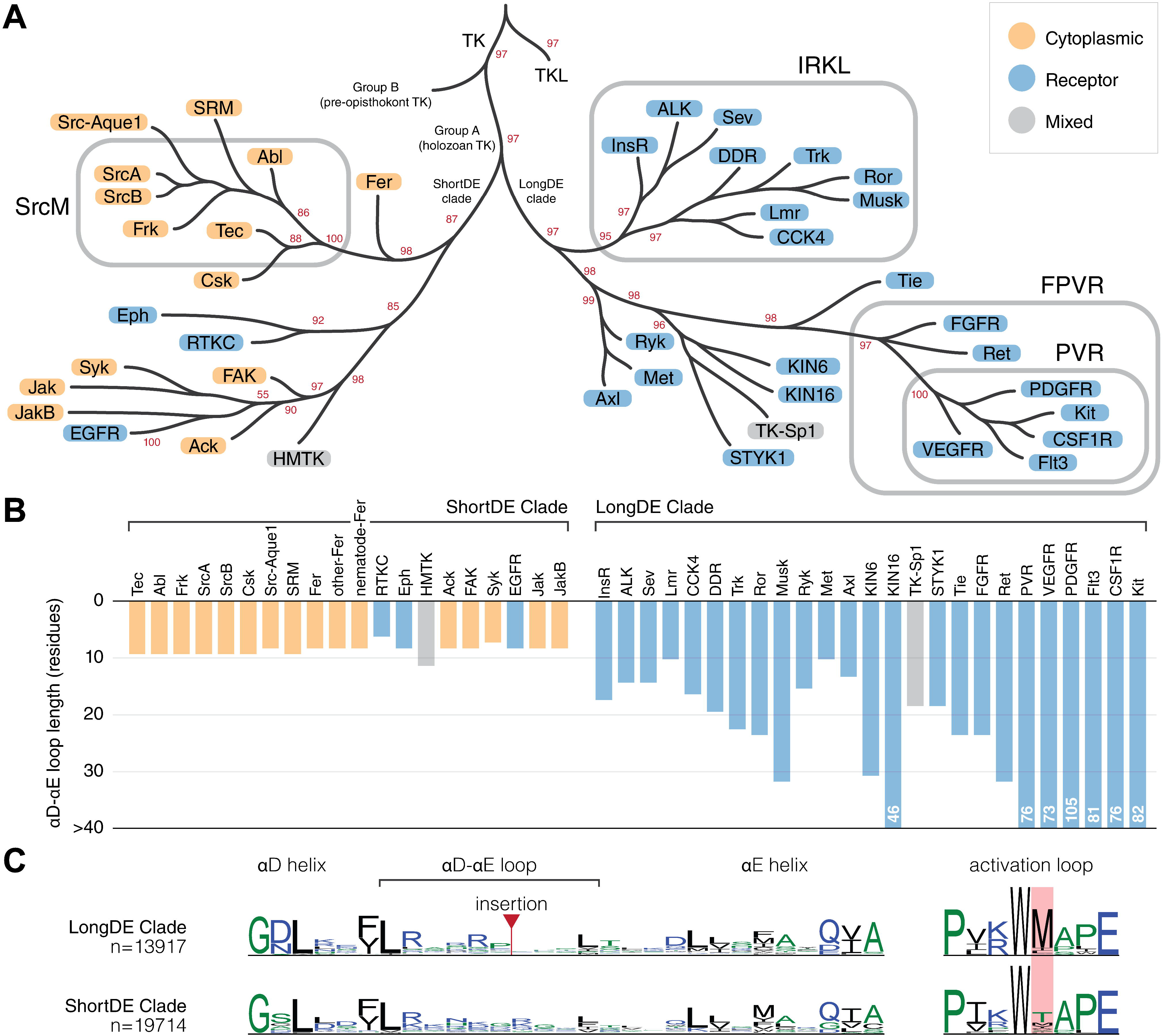
Evolution of sequence constraint-defined subgroups/families in the holozoan tyrosine kinome. (A) An abridged depiction of our representative phylogeny for the holozoan tyrosine kinome. Branch tips represent clades of sequence constraint-defined families. Cytoplasmic tyrosine kinase families are indicated with orange circles, while receptor tyrosine kinase families are indicated with blue circles. Subgroups are indicated by rounded rectangles which encompass their respective constituents. Bootstrap support values for select clades are shown in red. Branch lengths are not drawn to scale. The full tree is provided in **Supplemental File S4**. (B) A bar chart showing the median αD-αE loop length of each tyrosine kinase family, which is shown on the x-axis separated by LongDE or ShortDE clade. On the y-axis, loop length is truncated at 30 residues. Any αD-αE loop lengths surpassing this limit are designated as >30. (C) Comparative sequence logos show differences between ShortDE and LongDE clade kinases.

A further examination of our phylogenetic tree of tyrosine kinases reveals several new insights about tyrosine kinase evolution. As noted in previous phylogenetic studies of tyrosine kinase sequences, tyrosine kinases form a monophyletic clade separate from the closely related tyrosine kinase-like (TKL) group of serine/threonine kinases. Holozoan tyrosine kinases (Group A) also form a monophyletic clade that is distinct from a paraphyletic group of divergent tyrosine kinase sequences found in pre-opisthokonts (Group B), such as those found in the amoebozoans *Dictyostelium discoideum* or in the green algae *Chlamydomonas reinhardtii* **(Figure 6)** (Suga et al. 2012). We note for the first time another major branching point in the evolution of tyrosine kinases which is associated with the presence or absence of an insert between the αD and αE helices of the kinase domain that we refer to as the DE insert, historically referred to as the kinase insert domain (Locascio and Donoghue 2013). Although the DE insert segment was not explicitly used in building the phylogenetic tree due to lack of detectable sequence similarity in this region across families, the phylogenetic tree exhibits a clear division of two clades, one containing kinases with a short αD-αE loop, which we refer to as the shortDE clade, and one predominantly containing kinases with a long αD-αE loop, which we refer to as the longDE clade **(Figure 6A)**. The evolutionary separation of the longDE clade is also independently supported by a common phase-2 intron at the αH helix **(Supplementary Figure S6)**. While the variation in the length of the DE insert has been previously noted **(Figure 6B)**, and previous studies have shown the functional significance of the insert on downstream signaling and kinase activation (Locascio and Donoghue 2013; Manni et al. 2014), the evolutionary history of the DE insert has not been examined.

In light of the evolutionary divergence between longDE and shortDE clades, we also note that the longDE clade primarily contains receptor tyrosine kinase families, while the shortDE clade contains predominantly cytoplasmic receptor tyrosine kinases (with the exception of the EGFR, Eph, and choanoflagellate-specific RTKC families). This correlation between the presence of the longDE insert with the presence of transmembrane and extracellular domains, alongside evidence that the insert plays important roles in kinase activation and protein recruitment for downstream signaling, suggests that the longDE insert evolved as a means to facilitate downstream intracellular signaling upon the activation of receptor tyrosine kinases by extracellular signals. In addition, though the DE insert is difficult to align across families due to the lack of sequence conservation, the DE insert is alignable within families and often conserves sequence motifs including phosphorylatable tyrosine, serine, or threonine residues, suggesting that individual receptor tyrosine kinase families along the longDE clade have rapidly and frequently evolved the longDE insert in family-specific contexts, presumably to carry out family-specific downstream signaling functions. We also note that the longDE tyrosine kinases highly conserve a unique activation loop methionine, which is not observed in shortDE tyrosine kinases **(Figure 6C)**; however, the role of this methionine, or whether it is significant for DE insert function or for receptor tyrosine kinase function is unknown.

Interestingly, the shortDE clade in the tyrosine kinase phylogeny, which predominantly consists of cytoplasmic tyrosine kinases, includes two monophyletic clades of receptor tyrosine kinases: the EGFR family of receptor tyrosine kinases and a separate monophyletic clade that includes the Eph and choanoflagellate-specific RTKC families of receptor tyrosine kinases. Thus, our phylogeny suggests at least three independent origins of highly expanded receptor tyrosine clades, with the majority of receptor tyrosine kinases emerging from the longDE clade. That these disparate branches along the tyrosine kinase phylogeny have convergently evolved to include transmembrane and extracellular domains highlights the importance of relaying extracellular signals into intracellular responses across various signaling niches. While the longDE receptor tyrosine kinases are distinguished by extra functionalities imparted by the longDE insert, the Eph and EGFR families also exhibit unusual signaling functions so far unobserved in other longDE receptor tyrosine kinases. Eph receptor tyrosine kinases have a unique capacity for bi-directional signaling, where the binding of ephrin ligands, which are also membrane bound, can activate signaling both in the receptor-bearing cell, as well as in the ligand-bearing cell (Pasquale 2010). Furthermore, our tree suggests that the RTKC family may also share this unique capacity for bi-directional signaling. The EGFR family is also a unique family of receptor tyrosine kinases in that ligand binding induces dramatic conformational changes in the dimerization arm extracellularly, also inducing a unique allosterically activating dimer in the intracellular portion (Zhang et al. 2006; Jura et al. 2009). These mechanisms of EGFR family kinases, as well as their activating (rather than autoinhibitory) juxtamembrane and their lack of activation via activation loop phosphorylation distinguishes the EGFR family from receptor tyrosine kinases in the longDE clade (Lemmon and Schlessinger 2010; Lemmon et al. 2014).

## Discussion

The classification of protein kinases into evolutionarily related families has provided the foundation for decades of comparative sequence-structure-function studies on protein kinases (Hanks and Hunter 1995; Gerard Manning et al. 2002; G. Manning et al. 2002; Kwon et al. 2019). Here, we propose a new constraint-based classification of tyrosine kinases that newly defines the SrcM, IRKL, and FPVR subgroups **(Figure 2)** each of which maintains core subgroup-specific sequence motifs associated with subgroup-specific auto-inhibited conformations **(Figure 3)**. Subsequent taxonomic conservation analysis suggests that expansion of tyrosine kinase subgroups and evolution of family-specific motifs within these subgroups, along with domain shuffling, elaborated on subgroup-specific functions to diversify cell signaling functions **(Figure 4, 5)**. However, these core regulatory motifs are likely conserved because they ensure that these kinases are activated at the right place and time. For instance, the strongest SrcM-specific constraint is a methionine in the αC-β4 loop which packs against the auto-inhibited activation loop conformation, suggesting an anciently conserved role in stabilizing the Src-like inactive conformation **(Figure 3A)** that may be relieved in various manners such as the binding of substrates to the co-evolved SH3-SH2 domains. Furthermore, the ancient conservation of Csk and SrcM-specific constraints supports the pre-metazoan origins of SrcM inhibition via C-terminal tail phosphorylation by Csk (Taskinen et al. 2017). Further study of family-specific motifs across various lineages of tyrosine kinases is expected to reveal unique regulatory mechanisms across distinct tyrosine kinase families.

We constructed a new representative phylogenetic tree of the holozoan tyrosine kinome which revealed larger evolutionarily-related clades of tyrosine kinases associated with additional defining features **(Figure 6A)**. Dividing the holozoan tyrosine kinome into two roughly equal halves, the basal longDE and shortDE clades are distinguished by the presence or absence (respectively) of a fast-evolving kinase domain insertion in the αD-αE loop **(Figure 6B)**. Functionally important insertion regions shared by large groups of evolutionarily-related protein kinases have also been described in the CMGC group which conserves a kinase domain insertion between the αH and αI helices (Kannan and Neuwald 2004). The diverse functions of the longDE insertion remains understudied; however, the region is documented to possess many functionally-important phosphorylation sites in multiple tyrosine kinase families (Locascio and Donoghue 2013). Our tree also reveals three distinct evolutionary lines of receptor tyrosine kinases: longDE, RTKC-Eph, and EGFR, each of which are distinguished by unique lineage-specific variations on receptor tyrosine kinase signaling and regulation (Pasquale 2010; Locascio and Donoghue 2013; Lemmon et al. 2014). Overall, the definition of these evolutionarily-related tyrosine kinases enable researchers to infer sequence-structure-function relationships in the understudied kinases based on the known functions of well-studied kinase families.

The expansion and diversification of the tyrosine kinome across the animal kingdom highlights its central role in metazoan biology. While many previous studies have speculated on the role of tyrosine kinases in the evolution of multicellularity (Miller 2012; Hunter and Manning 2015; Brunet et al. 2017), our findings suggest key evolutionary innovations which likely contributed to the adoption of tyrosine kinase signaling for multicellular functions **(Figure 7A)**. While elaborate tyrosine kinase signaling networks have been discovered in unicellular pre-metazoans, they generally display low orthology to tyrosine signaling networks in metazoans (Manning et al. 2008). Our analyses identify sparse similarities between pre-metazoan and metazoan tyrosine kinase signaling in that SrcM, Tec, Csk, and IRKL tyrosine kinases originated in pre-metazoans and have remained conserved throughout diverse metazoan taxa **(Figure 4)**. These components may represent a core phospho-tyrosine signaling machinery that have been expanded through the addition of taxa-specific tyrosine kinases to suit unique organismal needs. We further speculate that additional capacity for bi-directional signaling emerged in the RTKC-Eph lineage in an extant ancestral pre-metazoan organism, as previous studies have identified distant Eph orthologs in various pre-metazoan organisms (Richter and King 2013; Tong et al. 2017) which are not shown in our conservation table because they lack Eph-specific sequence constraints. These core signaling functions likely enabled the evolution of primordial metazoa, a mass of cells capable of moving and growing in a coordinated fashion, lacking much of the complexity found in modern day metazoans. Throughout the course of metazoan evolution, the emergence of new complex biological systems such as the nervous system, circulatory system, and adaptive immunity appears to coincide with the emergence of functionally associated tyrosine kinase families **(Table 1)**. Additional complexity was gained through whole genome duplication and small-scale duplication events which played a major role in tyrosine kinase evolution in vertebrates, especially the PVR kinases whose evolution has been heavily influenced by both types of duplication event (Grassot et al. 2006; D’Aniello et al. 2008; Brunet et al. 2016; Brunet et al. 2017).

**Figure 7:**
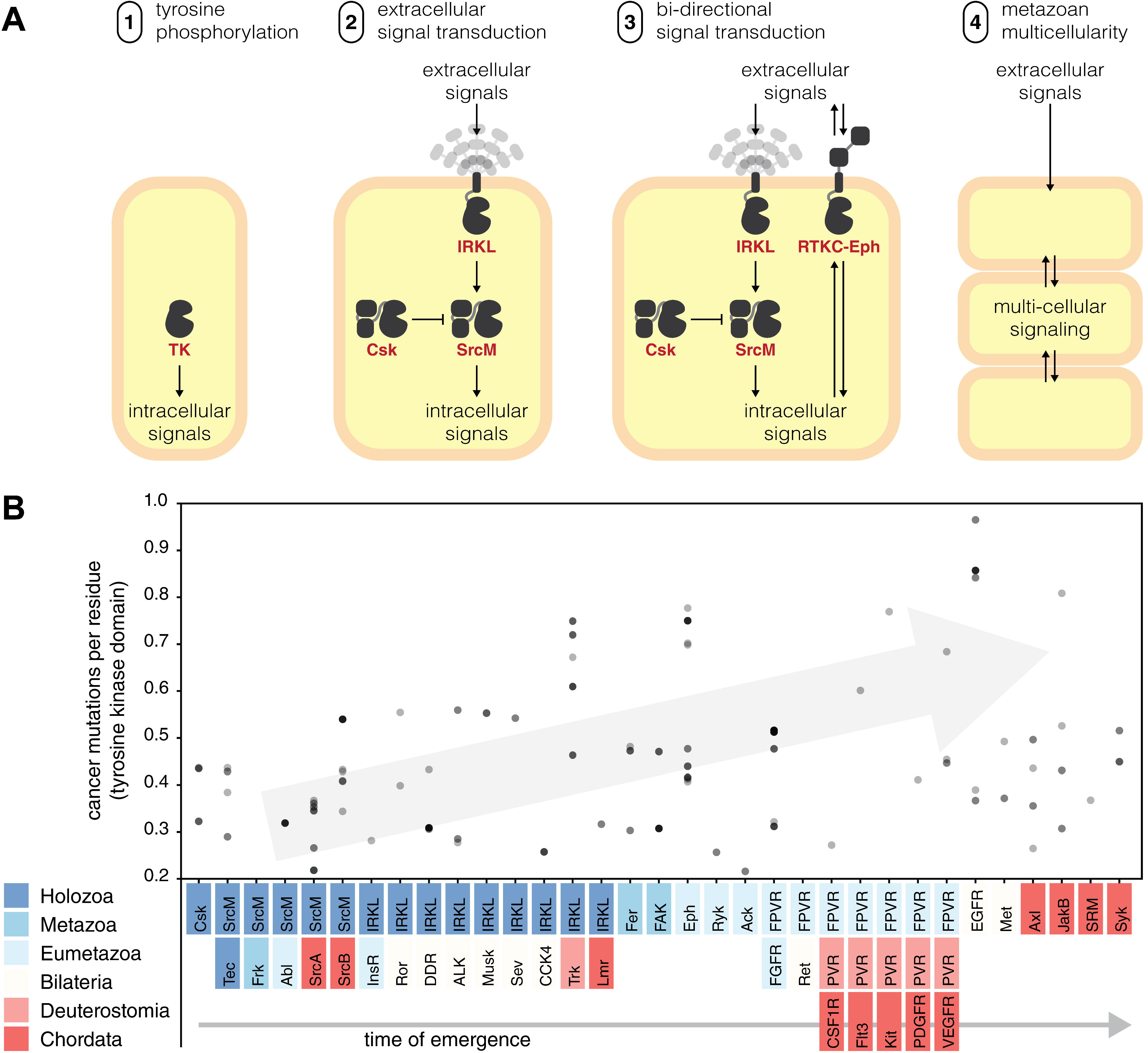
Tyrosine kinases in the evolution of multicellularity and cancer. (A) We propose a series of evolutionary innovations of the tyrosine kinome signaling which potentially contributed to the emergence of metazoan multicellularity. (B) Disease-related mutations tend to occur in more recent tyrosine kinase families. A scatter plot shows how frequently human tyrosine kinases are found to be mutated in genome-wide cancer sequencing studies from the Catalogue Of Somatic Mutations In Cancer (COSMIC) database (Tate et al. 2019). On the y-axis, mutation frequency is measured by the average number of mutations per residue in the tyrosine kinase domain. On the x-axis, each kinase is sorted into a cluster defined by our hierarchy and sorted by their time of emergence. All points were drawn with transparency; as a result, points which appear darker indicate multiple points falling in the same location.

**Table 1:** The emergence of tyrosine kinase families is associated with the metazoan phenotypes. The taxa in which many tyrosine kinase families first emerged **(Figure 4)** is correlated with the emergence of various taxa-specific phenotypes which are associated with functions performed by the corresponding tyrosine kinase family. Families whose functions do not seem to exhibit a clear connection to the emergence of a metazoan phenotype are not shown.

| family / subgroup | kinase functions | time of emergence | predicted association to metazoan evolution | references |
| --- | --- | --- | --- | --- |
| FAK | dynamically regulation of cell adhesion & motility | Porifera (Metazoa) | organized cell layer (found in all metazoans) | (Mitra et al. 2005; Mitchell and Nichols 2019) |
| Eph | regulation of cell migration during development | Cnidaria (Eumetazoa) | organized germ layers (emerged in cnidarians) | (Poliakov et al. 2004) |
| FGFR | regulation of early metazoan development, such as gastrulation and neural induction | Cnidaria (Eumetazoa) | simple nervous system (emerged in eumetazoans) | (Green et al. 1996; Matus et al. 2007; Bertrand et al. 2014) |
| FPVR | many FPVR kinases play roles in vasculature-related functions | Cnidaria (Eumetazoa) | vasculature (emerged in bilaterians) | (Gerard Manning et al. 2002) |
| EGFR | regulated cell growth via the EGFR pathway | Bilateria | EGFR pathway (emerged in bilaterians) | (Barberán et al. 2016) |
| Ror, Musk, DDR, Sev, ALK, CCK4 | many IRKL kinase families play roles in neural development | Bilateria | true brain characterized by neuropile (emerged in bilaterians) | (Gerard Manning et al. 2002; Chiu and Cline 2010) |
| VEGFR, PDGFR, Kit, CSF1R, Flt3 | many PVR kinase families play roles in blood vasculature-related functions and are implicated in blood cancers | Chordata | closed circulatory system lined with endothelium (emerged in vertebrates) | (Monahan-Earley et al. 2013; Berenstein 2015) |
| JAK, JAK-b, Syk | immune signaling via the JAK-STAT pathway | Chordata | adaptive immunity (emerged in vertebrates) | (Mócsai et al. 2010; Liongue et al. 2016) |

Given the important roles of tyrosine kinases in multicellular metazoan biology, it comes as no surprise that sequencing efforts have identified many disease-related variants across the tyrosine kinome (Hunter 2009). Multiple studies have proposed reversal of cancer cells to a more unicellular-like state through the release of “multicellular constraints” (Chen et al. 2015). While many oncogenes predate the origin of multicellularity, evolutionary studies have identified a second burst of oncogene emergence which co-occurred with the appearance of multicellular metazoans, and is comprised of genes (including tyrosine kinases) involved in cellular signaling and growth processes (Domazet-Lošo and Tautz 2010). In support of this finding, our data offers a closer look at this event, suggesting that cancer variants occur more frequently in recently-evolved tyrosine kinases **(Figure 7B)**. Overall, the deep connection between phospho-tyrosine signaling and cancer encourages further studies on the tyrosine kinome and the unique sequence constraints that guide their evolution and function. Focused analyses are expected to reveal new insights on metazoan multicellular signaling and its malfunctions in disease states.

## Materials and Methods

### Evolutionary constraint-based clustering

We sampled alternative classification hierarchies *ab initio*, using the omcBPPS algorithm (Neuwald 2014) which employs a Markov Chain Monte Carlo (MCMC) sampling strategy to classify aligned sequences into hierarchical categories/clusters based on shared constraints i.e. slow evolving sites. Category/cluster-specific constraint refers to alignment positions that are highly conserved within a given cluster, but divergent in sequences outside of the cluster. The omcBPPS algorithm iteratively optimizes for two interdependent criteria: (1) which alignment positions should be defined as cluster-specific constraints and (2) what is the optimal hierarchical clustering scheme based on cluster-specific constraints.

We ran omcBPPS on two different sequence sets from UniProt reference proteomes (retrieved 2/13/19) and NCBI non-redundant (nr) proteins database (retrieved 2/13/19) (Pruitt et al. 2007). Within these sequence sets, we identified and aligned tyrosine kinase sequences using the Multiply-Aligned Profiles for Global Alignment of Protein Sequences (MAPGAPS) algorithm (Neuwald 2009). This alignment was restricted to the protein kinase domain starting from the β1 strand (PKA^Phe43^) and ending at the αI helix (PKA^Lys292^). Kinase sequences which did not span from at least the β3-Lys to the DFG-Asp were deemed fragmentary and removed from the alignment. The UniProt sequence set contained 12,137 tyrosine kinase sequences. The nr sequence set was further purged at 98% sequence identity and contained 17,071 tyrosine kinase sequences. We performed hierarchical clustering on both sequence sets using omcBPPS. For both sequence sets, we optimized the “minnats” parameters (minnat=1 and minnat=5) which changed the minimum log-likelihood required to form a cluster. All runs were performed twice. To create a consensus of the hierarchical classification schemes found by multiple runs of omcBPPS, we used the mcBPPS algorithm. Clusters which were consistently identified throughout multiple runs were refined using the mcBPPS algorithm (Neuwald 2011). We ran mcBPPS on a maximally diverse set of 33,769 sequences containing all tyrosine kinase detectable from nr to generate an optimal model that is consistent with the existing data.

### Fitting new sequences to a constraint-based clustering model

Using our consensus model, we developed quantitative means of evaluating how well any given sequence fit into each of the clusters defined by our model **(Supplementary Algorithm S1)**. Within our model, each cluster was associated with a list of constraints that dictated which amino acid(s) were likely to be found at a given alignment position. Furthermore, each constraint was associated with a log-likelihood score which described how specific a constraint was to its respective cluster. To score a sequence against a cluster, we added the log-likelihoods of all the constraints which were true for the query sequence. In order to make this score comparable across clusters, we divided this number by the total log-likelihood of all the cluster’s constraints. This resulted in a range from 0 to 1. For example, a sequence which followed all of a given cluster’s constraints would have received a maximum score of 1 for that cluster, while a sequence which did not follow any of a given cluster’s constraints would have received a minimum score of 0 for that cluster. For the purposes of discrete classification, we defined a cutoff score for classifying a sequence into a cluster. Based on the distribution of scores from all possible sequence-cluster pairs across multiple test datasets, we defined the optimal cutoff at the global minima of 0.7.

Using **(Supplementary Algorithm S1)**, we scored a representative set of sequences from UniProt proteomes (retrieved 4/2/20) and estimated the size of each cluster. A representative set of 44,639 tyrosine kinase sequences spanning 586 species were identified and aligned to a common alignment profile using the MAPGAPS algorithm (Neuwald 2009). We determined the size of each cluster based on how many sequences scored above the cutoff. We also quantified the similarity between clusters by evaluating how well sequences in a given cluster scored against all other clusters using an all-vs-all comparison **(Supplementary Figure S2)**. We defined a non-symmetric similarity metric for any two given clusters, A and B, as the average score of cluster A sequences when classified against cluster B constraints. As a consequence of our cutoff, the average score when A and B were the same cluster was always greater than 0.7 which we observe across the diagonal. Indicative of our hierarchical classification scheme, we also observed a distinct signature for subclusters (such as Tec) which were classified under a larger supercluster (such as SrcM): subclusters would score highly (>0.7) for the constraints of its respective supercluster but not vice versa. Our comparison matrix also indicated that evolutionarily-related sequences within clusters share more constraints in common than sequences outside of the cluster. Finally, we observed that the majority of values outside of the diagonal were quite low which indicated that our model defined well-separated sequence clusters, each defined by their own unique sequence constraints.

### Comparative sequence analysis

All comparative sequence analyses were performed in Python 3 using the HelperBunny library (provided with our computational notebooks) which implements NumPy array-based sequence alignment manipulation. All sequence features (including features pertaining to motifs, evolutionary constraints, domain composition, taxonomic descriptors, insertions, and deletions) were represented as Boolean arrays as a function of the full sequence alignment. More complex queries pertaining to multiple sequence features (such as the presence of a given domain in a given taxon) were constructed by Boolean algebra. These Boolean arrays were applied as filters to our sequence alignment using NumPy indexing routines (Harris et al. 2020). Sequence logos were generated using the WebLogo 3 API (Crooks et al. 2004).

### Taxonomic conservation analysis

After we classified our representative set of tyrosine kinase into discrete clusters, we determined the taxonomic conservation of each cluster. We determined the source of each tyrosine kinase sequence using the organism identifier number (OX) provided in the FASTA header of UniProtKB sequences. OX numbers were traced back to their parent node identifiers using the nodes dump file provided in the NCBI taxdump database. All node identifiers were translated to their respective scientific names using the taxonomy names dump file. We determined the distribution of taxa across each cluster of tyrosine kinases and selected an optimal mix to depict diverse taxa ranging from unicellular pre-metazoans to more complex metazoans such as chordates **(Supplementary File S2)**.

### Intron and phase position analysis

Intron/exon annotations were mapped using Scipio (Keller et al. 2008). Intron phases were determined by calculating the modulus of three (representing the codon size) of the cumulative sum of lengths of each exon which preceded an intron within an open reading frame. A phase-0 intron is located between two consecutive codons; a phase-1 intron is located between the first and second nucleotides of a codon; and a phase-2 intron is located between the second and third nucleotides of a codon.

### Protein domain conservation analysis

We produced protein domain annotations for each full-length tyrosine kinase sequence using the NCBI Conserved Domain Database (CDD v3.18 - 55,570 PSSMs) database of conserved protein domains (Marchler-Bauer et al. 2011; Marchler-Bauer et al. 2017). Queries to the database were programmatically submitted using the bwrpsb PERL script which was provided in the Batch CD-Search API. Search parameters included an expected value threshold of 0.01 with only the best scoring domain model being returned. Transmembrane (TM) helix annotations were identified and appended to our domain annotation results using TMHMM 2.0 (Krogh et al. 2001). Synonymous domain names were manually identified and merged in post-processing. We combined these domain annotations with previously generated data to evaluate the conservation of protein domains across various constraint-defined clusters and taxonomic clades.

### Phylogenetic analysis

We inferred the evolution of tyrosine kinase families/subgroups using a maximum-likelihood approach. In order to create a representative phylogeny of the holozoan tyrosine kinome, we sampled a taxonomically diverse set of sequences from each constraint-defined cluster of tyrosine kinase sequences. Our representative set of sequences consisted of (1) one randomly selected sequence from each cluster-taxon pair (based on **Supplementary File S2**) excluding chordata, (2) all human tyrosine kinases sequences which represented the chordate taxon, (3) three early-diverging, pre-opisthokont tyrosine kinase sequences from amoebozoan, *A. castellanii* (Suga et al. 2012), and (4) eight human TKL group kinases which was used as an outgroup. We produced multiple representative sequence sets with different random samples of taxonomically diverse sequences. Using these sequence sets, we generated multiple phylogenies using IQTREE v1.6.11 (Nguyen et al. 2015), with ModelFinder (Kalyaanamoorthy et al. 2017). Branch support values were generated using ultrafast bootstrap with 1000 resamples (Hoang et al. 2018). Results indicated that sequences from the same clusters consistently formed paraphyletic groups or monophyletic clades with high bootstrap support **(Supplementary File S4, S5)**. We compared topologies generated from different random samples and found no major changes in the evolutionary relationships between evolutionary constraint-defined sequence clusters. Furthermore, our topology was robust to the inclusion of unclassified tyrosine kinase sequences.

The consensus tree with the highest bootstrap support values for the three major tyrosine subgroups was selected as the final tree. The optimal substitution model for our final topology was determined to be LG+R8 based on the Bayesian Information Criterion as determined by ModelFinder (Kalyaanamoorthy et al. 2017). We rooted our final tree against the TKL outgroup using ETE Toolkit v3.1.1 (Huerta-Cepas et al. 2016). Consistent with previous studies, the most divergent tyrosine kinases in our tree were the paraphyletic pre-opisthokont “Group B” tyrosine kinases which diverged prior to the emergence of the monophyletic “Group A” holozoan tyrosine kinases (Suga et al. 2012). We observed high concordance between our evolutionary constraint-based clustering and phylogenetic inference; this allowed us to simplify our tree topology by condensing monophyletic clades and pruning paraphyletic groups **(Figure 6A)**. The simplified representation describes evolutionary relationships between constraint-defined families. Furthermore, we represented each of the three major tyrosine kinase subgroups by drawing an enclosure around each subgroup’s constituent families.

We compared our tree to previously published phylogenies which also sampled diverse tyrosine kinase families (Robinson et al. 2000; G. Manning et al. 2002; Suga et al. 2008; Suga et al. 2012; Brunet et al. 2016; Modi and Dunbrack 2019). However, many of these studies focused on vertebrate, basal metazoan, and pre-metazoan kinase sequences leaving out many diverse protostome sequences. We observed key differences with the current widely-accepted phylogeny of the human tyrosine kinome **(Supplementary Figure S3)** (G. Manning et al. 2002) and found the most similarities to a tree published by Robinson et al (Robinson et al. 2000). Previously placed near the base of the tyrosine kinase clade, our placement of STYK1 was supported by common introns shared with Tie and various IRKL families, while our placement of Lmr was supported by common introns shared by various IRKL kinases (Brunet et al. 2016; Brunet et al. 2017). Furthermore, the majority of human longDE kinases share a common phase-2 intron in the genomic region which codes for the αH helix **(Supplementary Figure S6)**. Overall, our tree has high support for all major clades, including historically difficult-to-place families such as JAK (Shiu and Li 2004). Providing additional support, we note that our tree is highly concordant with the evolutionary progression of holozoan taxa in that anciently conserved tyrosine kinase families tend to diverge first, while more recently diverged tyrosine kinase families appear closer to the tips **(Supplementary Figure S4)**.

## Supporting information

Supplemental Algorithm S1

## Acknowledgements and funding information

We thank Dr. James H Leebens-Mack for valuable feedback. This work was supported by the National Institutes of Health (R01 GM114409 and R35 GM139656 to NK).

## Data availability statement

The datasets and code for our analyses are freely available for download from GitHub at: https://github.com/esbgkannan/holozoan_tk_evolution.

