## Supplemental Text for "Evolution of functional diversity in the holozoan tyrosine kinome"

### Supplementary Figure Legends

**Supplemental Figure S1: A hierarchical, constraint -based classification of tyrosine kinase sequences.** The previously determined classification (left) and the new hierarchical, constraint-based classification (right) are depicted as cladograms. Each node represents a tyrosine kinase subgroup or family, with the number in parentheses indicating the number of sequences classified into each subgroup/family. Labels above each branch indicate log probability ratio (LPR) scores for each subtree in the classification (calculated by the mcBPPS algorithm). The total LPR scores are indicated above each cladogram.

**Supplemental Figure S2: Quantifying similarities between clusters of tyrosine kinases.** Each cluster of sequences (x-axis) is scored against the evolutionary constraints associated with each cluster (y-axis). The average score for each cluster-pattern set pair is shown on the heat map. Sequences are defined as being a member of a cluster if its pattern score is at least 0.7. This is reflected by the high scores along the diagonal. Clusters which contain sub-clusters are characterized by horizontal rows containing more than one value greater than 0.7. These super-clusters include PVR, IRKL, SrcM, and Fer.

**Supplemental Figure S3: Comparison of constraint-guided phylogeny and the Manning phylogeny of the human kinome.** (A) On the left, we show the tyrosine kinase group from Manning's phylogeny of the human tyrosine kinome. On the right, we collapse all clades containing each of the lowest level clusters defined by our constraint-based classification. Cytoplasmic tyrosine kinases are highlighted orange, while receptor tyrosine kinases are highlighted blue. (B) Our constraint-guided phylogeny (left) is compared to the Manning's phylogeny of the human tyrosine kinome (right). Matching lines are drawn to facilitate comparison. We use simplified versions of both phylogenies to facilitate this comparison. Our constraint-guided phylogeny was simplified as described in the main text. However, the multifurcation between FGFR, Ret, and PVR was resolved based on their taxonomic conservation depth to produce a bifurcating tree. Red clades denote clades which are not found in the Manning phylogeny. The Manning phylogeny was pruned as by cutting monophyletic clades described by our constraint-based clustering. The human kinases found in each pruned monophyletic clade is shown on the right side.

**Supplemental Figure S4: The representative phylogeny of the holozoan tyrosine kinome is highly concordant with the evolutionary progression of holozoan taxa.** A condensed phylogeny of the holozoan tyrosine kinome. Constraint-based tyrosine kinase subgroups are encircled, and subgroups and families are colored according to their inferred evolutionary time of emergence as indicated by the taxonomic clades labeled in the legend. Bootstrap support values for select clades are shown in red.

**Supplemental Figure S5: Lmr kinases have regained the serine/threonine kinase-specific histidine in the αH helix which was previously lost in tyrosine kinases.** Comparative sequence logos show differences between Lmr family kinases, tyrosine kinases, and serine/threonine kinases. The serine/threonine kinase-specific histidine is highlighted in red.

**Supplemental Figure S6: The representative phylogeny of the holozoan tyrosine kinome is highly concordant with common intron positions and phases found in humans.** We plot the intron positions and phases of all human tyrosine kinases and map their relative positions on the kinase domain using our alignment. Our representative phylogeny of the holozoan tyrosine kinome is shown across the y-axis. Conserved regions of the kinase domain are shown on top, across the x-axis. Each human tyrosine kinase domain is represented by a horizontal grey line with colored ticks which indicate intron positions and phases. As indicated by the legend (top-left), phase-0 introns are represented by a light blue tick, phase-1 introns are represented by a crimson tick, and phase-2 introns are represented by a maroon tick.

### Supplementary Algorithms

**Supplemental Algorithm S1:** Quantification of sequence-cluster fit.

### Supplementary Files

**Supplemental File S1: Statistics for constraint-defined tyrosine kinases clusters.** We list statistics for all constraint-defined subgroups/families in our analysis. Receptor tyrosine kinase (RTK) definitions are based on whether the kinase has a transmembrane helix detected by TMHMM (Krogh et al. 2001). Pseudokinases were defined are based on whether the kinase co-conserves the β3-Lys, HRD-Asp, and DFG-Asp (Kwon et al. 2019).

**Supplemental File S2: Taxonomic conservation of tyrosine kinases families across diverse holozoan taxa.** We detect the presence of evolutionary constraints throughout tyrosine kinases within holozoans (animals and related single-celled organisms excluding fungi). Tyrosine kinase subgroups/families are shown across the x axis, while holozoan taxa are shown across the y axis. The parenthetical numbers denote the species represented within each taxon. The unclassified category includes tyrosine kinases which did not fit into any clusters defined by our model.

**Supplemental File S3: Comparison of classification schemes for human tyrosine kinases.** A table of 94 human tyrosine kinase domains is shown. We provide the hierarchical classification of each kinase using our evolutionary constraint-based classification compared to the KinBase classification.

**Supplemental File S4: Representative phylogeny of the holozoan tyrosine kinome.** We construct a phylogenetic tree using kinase domain sequences from all human tyrosine kinases and a representative sample of tyrosine kinases from diverse organisms. The phylogeny was rooted to an outgroup of 8 TKL kinases. Three early-diverging, pre-opisthokont tyrosine kinases (Suga et al. 2012) are included as reference. Kinases belonging to one of the three major subgroups are highlighted: SrcM (blue), IRKL (gold), and FPVR (green).

**Supplemental File S5:** Newick tree for the representative phylogeny of the holozoan tyrosine kinome.

**Supplemental File S6:** Sequence alignment for the representative phylogeny of the holozoan tyrosine kinome.

### Supplementary Figures

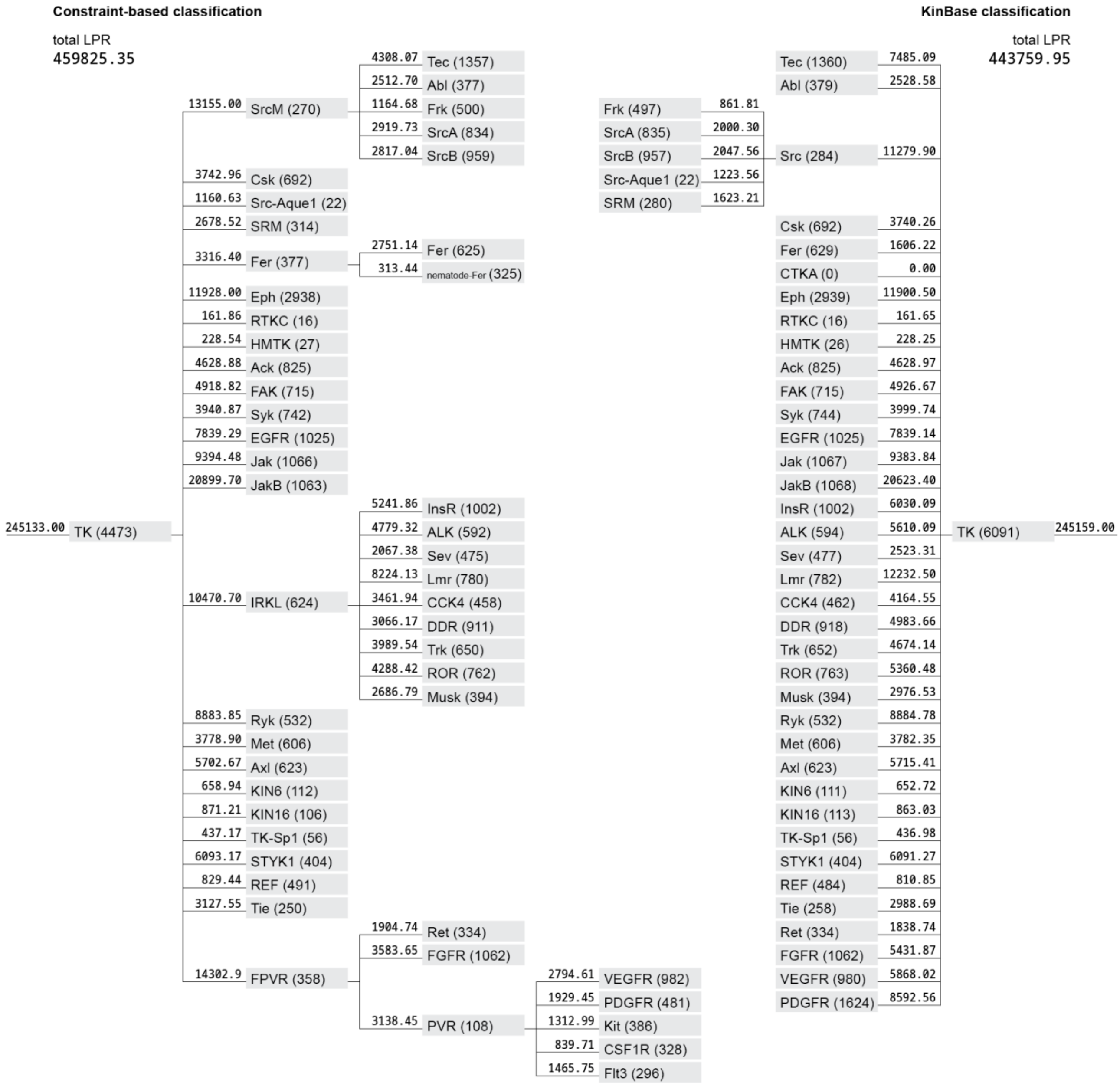


**Supplemental Figure S1: A hierarchical, constraint-based classification of tyrosine kinase sequences.**

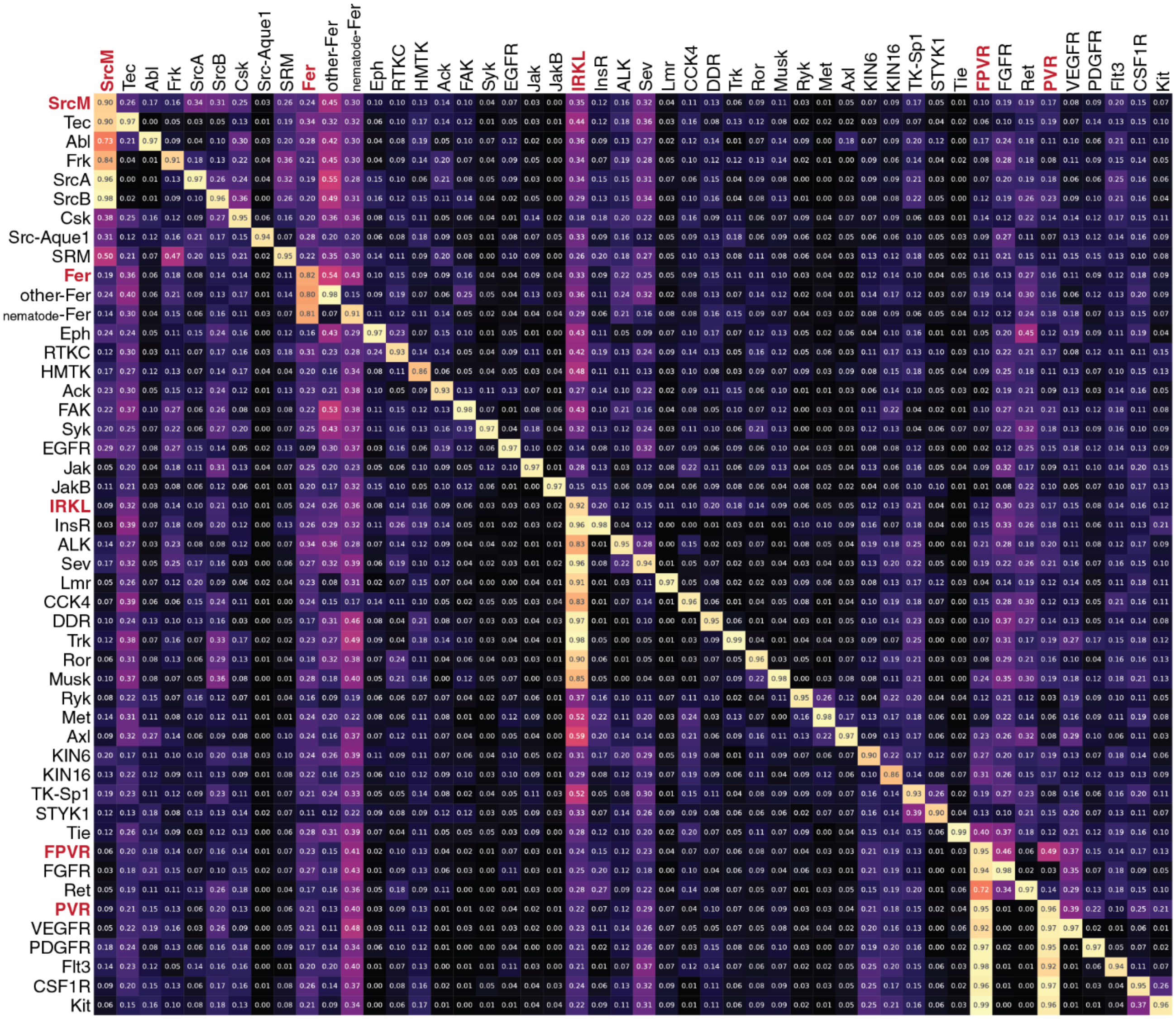


**Supplemental Figure S2: Quantifying similarities between clusters of tyrosine kinases.**

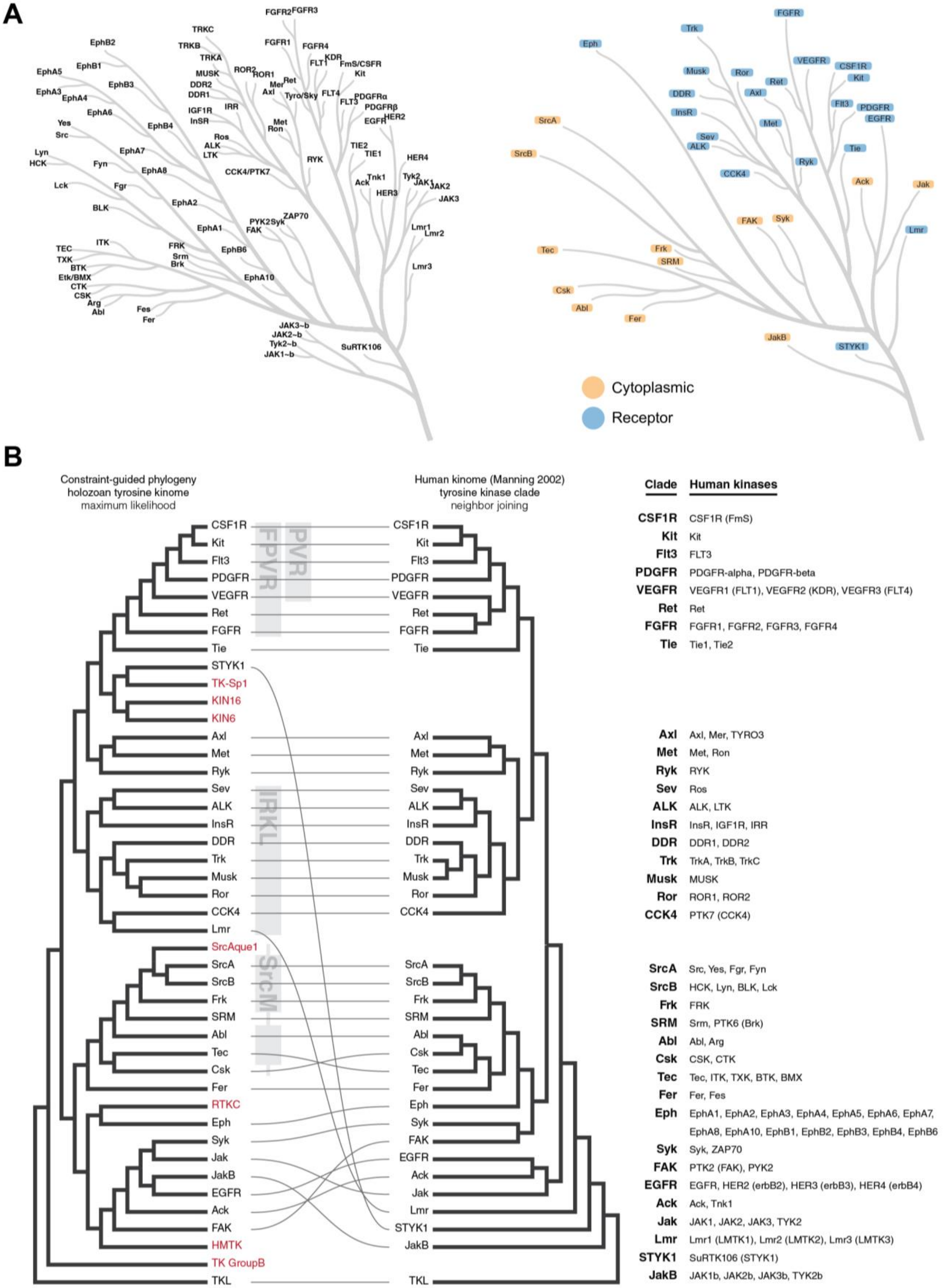


**Supplemental Figure S3: Comparison of constraint-guided phylogeny and the Manning phylogeny of the human kinome.**

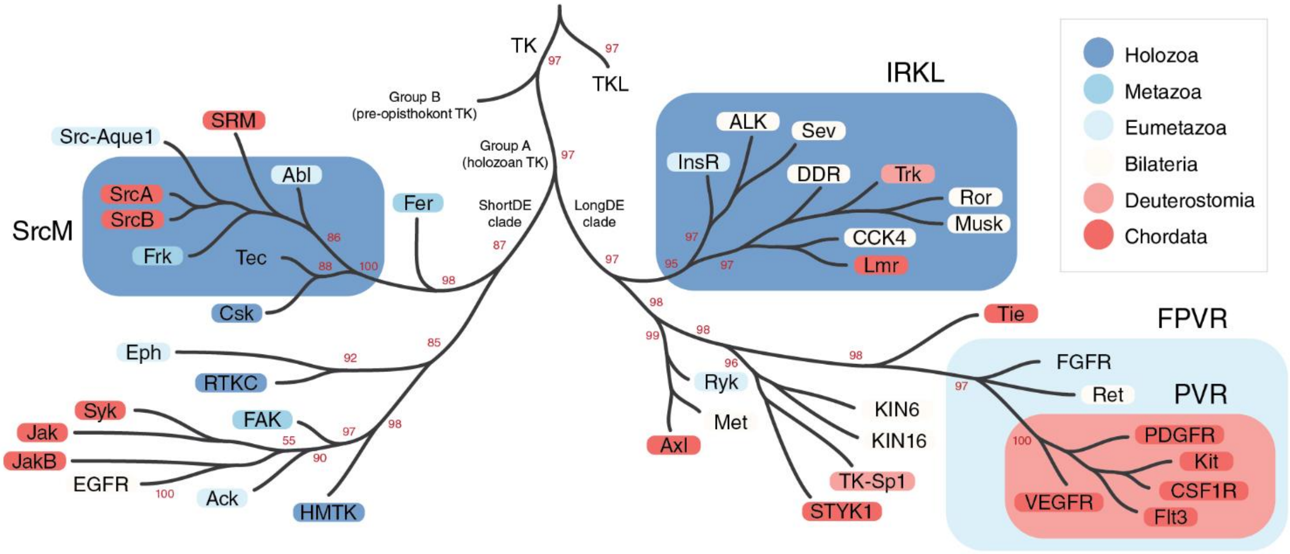


**Supplemental Figure S4: The representative phylogeny of the holozoan tyrosine kinome is highly concordant with the evolutionary progression of holozoan taxa.**

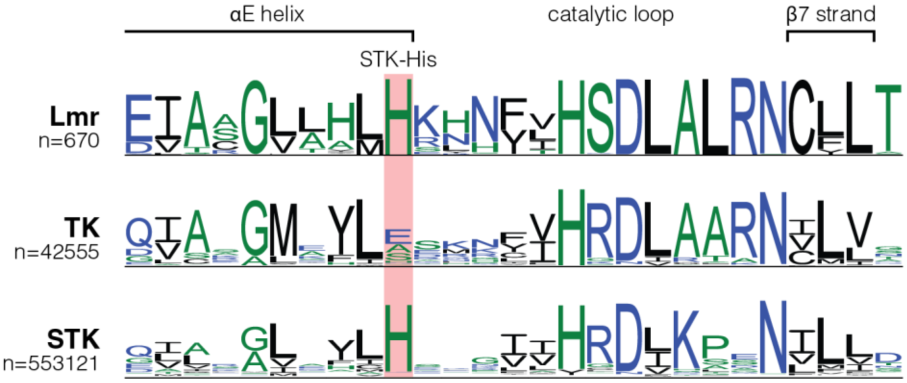


**Supplemental Figure S5: Lmr kinases have regained the serine/threonine kinase-specific histidine in the αH helix which was previously lost in tyrosine kinases.**

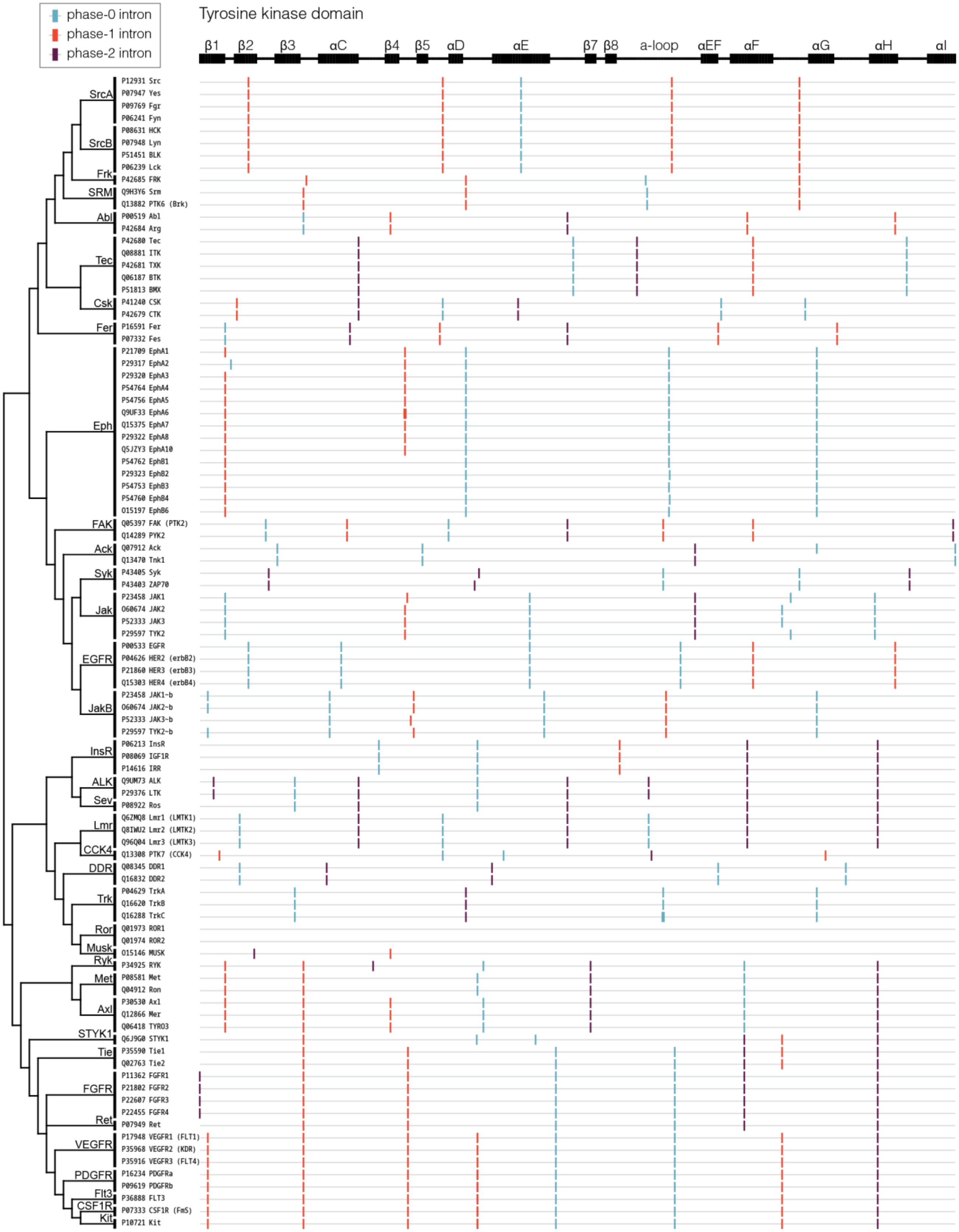


**Supplemental Figure S6: The representative phylogeny of the holozoan tyrosine kinome is highly concordant with common intron positions and phases found in humans.**
