## Supplemental Algorithm S1 for "Evolution of functional diversity in the holozoan tyrosine kinome"

**Algorithm:** Quantification of sequence-cluster fit

**Definitions:**

$$\begin{aligned} S &= \text{Query Sequence} \\ C_{cluster} &= \text{All Constraints for a given Cluster} \\ c &= \text{Constraint} \\ S_L &= \text{Sum of Log likelihoods for Query Sequence} \\ c_L &= \text{Log likelihood for a Constraint} \\ Total &= \text{Sum of Log likelihoods for a given Cluster} \end{aligned}$$

**for** $c$ ***in*** $C_{cluster}$ **do**

$$S_L \mathrel{+}= \begin{cases} c_L, & \textit{if c is true for S} \\ 0, & \textit{otherwise} \end{cases} \tag{1}$$

$$Total \mathrel{+}= c_L \tag{2}$$

**end**

$$Score = \frac{S_L}{Total} \tag{3}$$
