## Supplementary figures and images for "Evolution of functional diversity in the holozoan tyrosine kinome"

### Supplemental File S4

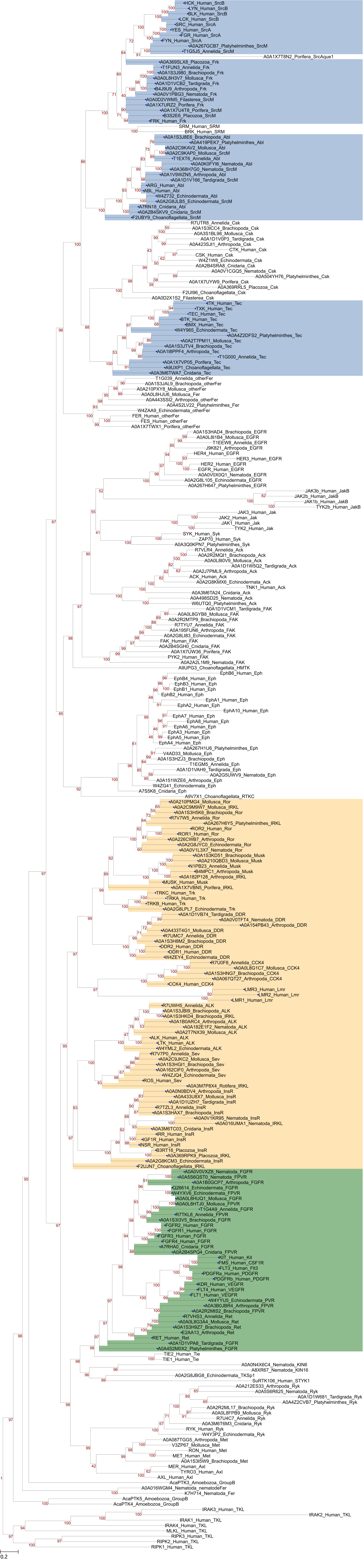
